# Adenosine Reduces Sinoatrial Node Cell AP Firing Rate by uncoupling its Membrane and Calcium Clocks

**DOI:** 10.1101/2022.06.20.496874

**Authors:** Ashley N. Wirth, Kenta Tsutsui, Victor A. Maltsev, Edward G. Lakatta

## Abstract

The spontaneous action potential (AP) firing rate of sinoatrial nodal cells (SANC) is regulated by a system of intracellular Ca^2+^ and membrane ion current clocks driven by Ca^2+^-calmodulin-activated adenylyl cyclase-protein kinase A (PKA) signaling. The mean AP cycle length (APCL) and APCL variability inform on the effectiveness of clock coupling. Endogenous ATP metabolite adenosine (ado) binds to adenosine receptors that couple to G_i_ protein-coupled receptors, reducing spontaneous AP firing rate via G_βγ_ signaling that activates an membrane-clock outward current, I_KACh_. Ado also inhibits adenylyl cyclase activity via G_iα_ signaling, impacting cAMP-mediated PKA-dependent protein phosphorylation and intracellular Ca^2+^ cycling. We hypothesize that in addition to I_KAdo_ activation, ado signaling impacts Ca^2+^ via G_iα_ signaling and that both effects reduce AP firing rate by reducing the effectiveness of the Ca^2+^ and membrane clock coupling. To this end, we measured Ca^2+^ and membrane potential characteristics in enzymatically isolated single rabbit SANC. 10 µM ado substantially increased both the mean APCL (on average by 43%, n=10) and AP beat-to-beat variability from 5.1±1.7% to 7.2±2.0% (n=10) measured via membrane potential and 5.0±2.2 to 10.6±5.9 (n=40) measured via Ca^2+^ (assessed as the coefficient of variability, CV=SD/mean). These effects were mediated by hyperpolarization of the maximum diastolic membrane potential (membrane clock effect) and suppression of diastolic spontaneous, local Ca^2+^ releases (LCRs) (Ca^2+^ clock effect): as LCR size distributions shifted from larger to smaller values, the time of LCR occurrence during diastolic depolarization (LCR period) became prolonged, and the ensemble LCR Ca^2+^ signal became reduced. The tight linear relationship of coupling between LCR period to the APCL in the presence of ado “drifted” upward and leftward, i.e. for a given LCR period, APCL was prolonged, becoming non-linear indicating clock uncoupling. An extreme case of uncoupling occurred at higher ado concentrations (>100 µM): small stochastic LCRs of the Ca^2+^ clock failed to self-organize and synchronize to the membrane clock, thus creating a failed attempt to generate an AP resulting in arrhythmia and cessation of AP firing. Thus, the effects of ado to activate G_βγ_ and I_KACh, Ado_ and to activate G_iα_, suppressing adenylyl cyclase activity, both contribute to the ado-induced increase in the mean APCL and APCL variability by reducing the fidelity of clock coupling and AP firing rate.

## Introduction

Cardiac sinoatrial nodal (SAN) pacemaker cells (SANC) generate spontaneous action potentials (APs) that initiate each heartbeat. Adenosine (ado), an endogenous cardiac metabolite, generated via the enzymatic hydrolysis of AMP or S-adenosyl homocysteine (Schrader, 1983;Shen and Kurachi, 1995). Ado concentration increases when ATP becomes reduced during metabolic stress and works to slow heart rate, thereby reducing energy consumption to protect the heart (West and Belardinelli, 1985;Belardinelli et al., 1988).

Ado has been previously shown to slow SAN impulses and delay SAN conduction (Drury and Szent-Gyorgyi, 1929). Applications of ado to single isolated SANC reduce the AP firing rate (Belardinelli et al., 1988;Headrick et al., 2011) via ado-activated A_1_ receptor signaling that leads to activation of inhibitory GTP-binding protein to dissociate into its G_iα_ and G_βγ_ subunits. These effects, respectively, result in the inhibition of adenyl cyclase (AC) and activation of I_KACh_ current which hyperpolarizes the SANC membrane potential (Logothetis et al., 1987;Belardinelli et al., 1988;Shen and Kurachi, 1995). It had been postulated (Belardinelli et al., 1988) and shown directly (Kurachi et al., 1986) that ado and Acetylcholine (Ach), which both activate G_βγ,_ target the same K^+^ channels (I_KACh,Ado_). The effects of ado and Ach to slow SANC AP firing rate have been, and continue to be, mainly attributed to this mechanism.

But more recently, evidence has emerged to indicate that SANC AP firing is regulated not solely by surface membrane ion currents, but by a coupled-oscillator system, driven by Ca^2+^-calmodulin activated AC-PKA and CaMKII signaling current (Lakatta et al., 2010). An intracellular Ca^2+^ oscillator or “Ca^2+^ clock”, the sarcoplasmic reticulum (SR), couples to an ensemble of voltage and time-dependent surface membrane current oscillators “Membrane clock”. The Ca^2+^ clock generates spontaneous, rhythmic local increases in diastolic Ca^2+^, local Ca^2+^ releases (LCRs), that activate inward Na^+^/Ca^2+^ exchanger current (I_NCX_), which partners with I_f_ to initiate activation of diastolic depolarization (Vinogradova et al., 2004). Feed-forward signaling involving low voltage activated Ca^2+^ channels (Ca_γ_1.3) (Torrente et al., 2016), Ca^2+^-induced Ca^2+^ release (CICR), and continued spontaneous LCRs from the SR form an ensemble Ca^2+^ signal that accelerates diastolic membrane depolarization via I_NCX_ activation (Lyashkov et al., 2018). When this self-organized electro-chemical oscillation achieves the membrane potential (V_m_) depolarized sufficient to open L-type Ca^2+^ channels, an AP is ignited, which (via CICR) induces a relatively synchronous activation of RyR, resulting in a global cytosolic Ca^2+^ transient (Fabiato, 1983;Stern et al., 1999). AP firing itself, via its effects to regulate intracellular Ca^2+^, the “oscillatory substrate” of the Ca^2+^ clock, affects Ca^2+^-ligand function of proteins of both clocks and thus affects clock coupling (Lyashkov et al., 2009;Maltsev and Lakatta, 2010). The average AP firing rate and AP cycle length (APCL) inform on the fidelity of clock coupling: when clock coupling decreases, the mean AP firing rate is reduced and the variability of APCL increases (Yaniv et al., 2014b;Moen et al., 2019).

Muscarinic cholinergic receptor activation via G_i_ protein activation directly activates I_KAch_ via G_βγ,_ to hyperpolarize the MDP (DiFrancesco and Tromba, 1988;Demir et al., 1999) and, via G_iα_ activation, inhibits AC activity, reducing cAMP-mediated, PKA-dependent protein phosphorylation (Dessauer et al., 1996;Jurevicius and Fischmeister, 1996;Cabrera-Vera et al., 2003;Okumura et al., 2003;Lyashkov et al., 2009). Accordingly, inhibition of cAMP-PKA signaling in response to cholinergic receptor stimulation has a direct effect to reduce intracellular Ca^2+^ cycling, reducing Ca-calmodulin activated AC-PKA-CAMKII (Lyashkov et al., 2009). This effect, in conjunction with the activation of K^+^ channels, reduces the mean AP firing rate (Lyashkov et al., 2009) and increases AP beat-beat variability (Yaniv et al., 2014a). Reduced AP firing reduces net Ca^2+^ influx and therefore intracellular Ca^2+^, the “oscillatory substrate” of the Ca^2+^ clock. Thus, Ach reduces functions of the clocks and the effectiveness of the clocks coupling via several intertwined Ca^2+^ and voltage-dependent mechanisms.

Effects of ado on intracellular Ca^2+^ cycling and Ca^2+^ and membrane clock coupling in SANC have not yet been evaluated in SANC. Because ado exerts its effects via a signaling cascade that involves the same G_i_ protein coupling as Ach that inhibits adenylyl cyclase activity via a G_iα_ effect and activates K^+^ channels via G_βγ_ effect, we hypothesize, that ado, like Ach, effects changes in both intracellular Ca^2+^ and V_m_ that reduces the membrane and Ca^2+^ clock functions and reduces clock coupling, culminating in AP firing rate reduction.

## Methods

### Single Cell Preparation

SANC were isolated from male rabbits in accordance with NIH guidelines for the care and use of animals, protocol # 34-LCS-2019 (as previously described) (Vinogradova et al., 2000). New Zealand White rabbits (Charles River Laboratories, USA) weighing 2.8–3.2 Kg were anesthetized with sodium pentobarbital (50–90 mg/kg). The heart was removed quickly and placed in solution containing (in mM): 130 NaCl, 24 NaHCO_3_, 1.2 NaH_2_PO_4_, 1.0 MgCl_2_, 1.8 CaCl_2_, 4.0 KCl, 5.6 glucose equilibrated with 95% O_2_ / 5% CO_2_ (pH 7.4 at 35°C). The SAN region was cut into small strips (∼1.0 mm wide) perpendicular to the crista terminalis and excised as reported previously (Vinogradova et al., 2000). The final SA node preparation, which consisted of SA node strips attached to the small portion of crista terminalis, was washed twice in Ca^2+^-free solution containing (in mM): 140 NaCl, 5.4 KCl, 0.5 MgCl2, 0.33 NaH_2_PO_4_, 5 HEPES, 5.5 glucose, (pH=6.9) and incubated on shaker at 35°C for 30 min in the same solution with the addition of elastase type IV (0.6 mg/ml; Sigma, Chemical Co.), collagenase type 2 (0.8 mg/ml; Worthington, NJ, USA), Protease XIV (0.12 mg/ml; Sigma, Chemical Co.), and 0.1% bovine serum albumin (Sigma, Chemical Co.). The SA node preparation was next placed in modified Kraftbruhe (KB) solution, containing (in mM): 70 potassium glutamate, 30 KCl, 10 KH_2_PO_4_, 1 MgCl_2_, 20 taurine, 10 glucose, 0.3 EGTA, and 10 HEPES (titrated to pH 7.4 with KOH), and kept at 4°C for 1h in KB solution containing 50 mg/ml polyvinylpyrrolidone (PVP, Sigma, Chemical Co.). Finally, cells were dispersed from the SA node preparation by gentle pipetting in the KB solution and stored at 4°C.

### High Speed 2D Ca^2+^ Signal Imaging

Ca^2+^ dynamics within isolated single human SANC were measured by 2D imaging of the fluorescent Ca^2+^ indicator, Fluo-4. Cells were loaded with 5 μM Fluo-4AM (Thermo Fisher, USA) for 15 minutes at room temperature. Fluo-4AM was subsequently washed out of the chamber with bathing solution contained the following (in mM): NaCl 140, HEPES 5, NaH_2_PO_4_*H2O 0.33, KCl 5.4, MgCl_2_ 1.0, glucose 5.5, CaCl_2_ 1.8; titrated to pH 7.35 with NaOH. Ca^2+^ signals were measured within the ensuing 30 minutes at 35°C ±0.1°C. Temperature was controlled by an Analog TC2BIP 2/3Ch bipolar temperature controller from CellMicroControls (USA), which heated both the glass bottom of the perfusion chamber and the solution entering the chamber (via a pre-heater).

Fluo-4 fluorescence was detected using a high-speed PCO.edge 4.2 CMOS camera (100 frames-second, with an 13.2 mm square sensor of 2048×2048 pixels resolution) mounted on a Zeiss inverted microscope (Carl Zeiss, Inc., Germany) with a x63 oil immersion lens and a fluorescence excitation light source (CoolLED pE-300-W, BioVision Technologies, Inc. PA, USA). Fluo-4 fluorescence excitation (blue light, 470/40 nm) and emission light collection (green light, 525/50 nm) were performed using the Zeiss filter set 38 HE. To avoid phototoxicity, Fluo-4 was excited only for short periods of time (<20 s) (Monfredi et al., 2013;Kim et al., 2018). Data acquisition was performed using PCO camware 64 (PCO AG, Germany).

### Membrane Potential Recording

V_m_ was measured in the current clamp configuration using an Axopatch 200B amplifier (Molecular Devices). Patch pipette resistances ranged between 3-5 MΩ, and pipettes were filled with a solution containing (in mM): K^+^ gluconate 120, NaCl 5, MgATP 5, HEPES 5, KCl 20; titrated to pH 7.2 with KOH. Amphotericin B (320 μM, Sigma-Aldrich A-4888) was added into the pipette solution as the pore-forming agent. Liquid junction potential was calculated by pClamp software (Molecular Devices) and adjusted accordingly.

### Computational Analysis of LCRs and APs

We used an in-house custom program (‘XYT Event Detector’) to objectively, automatically, and rapidly analyze the individual and ensemble behavior of the LCRs (Maltsev et al., 2017). The program yields detailed information about the number, timing, and size of individual LCRs. Specifically, The LCR timing is assessed as ‘LCR period’ which is the time interval between the peak of the prior AP-induced Ca^2+^ transient peak and the onset of the LCR occurrence. The “LCR size” is given as the LCR full propagation path in µm^2^. The computer program also provides the LCR ensemble signal (i.e. the summation of all LCR Ca^2+^ signal areas occurring within a given time, Fig. 1D). APs were analyzed by using another in-house custom program “AP analysis” (Lyashkov et al., 2018). In addition to traditional parameters APCL, APD50, MDP, and Take-Off potential, the program also provided AP ignition parameters introduced in (Lyashkov et al., 2018): Ignition duration (Idur), and Time-To-Ignition onset (TTI) (Fig. 1C).

**Figure 1.**
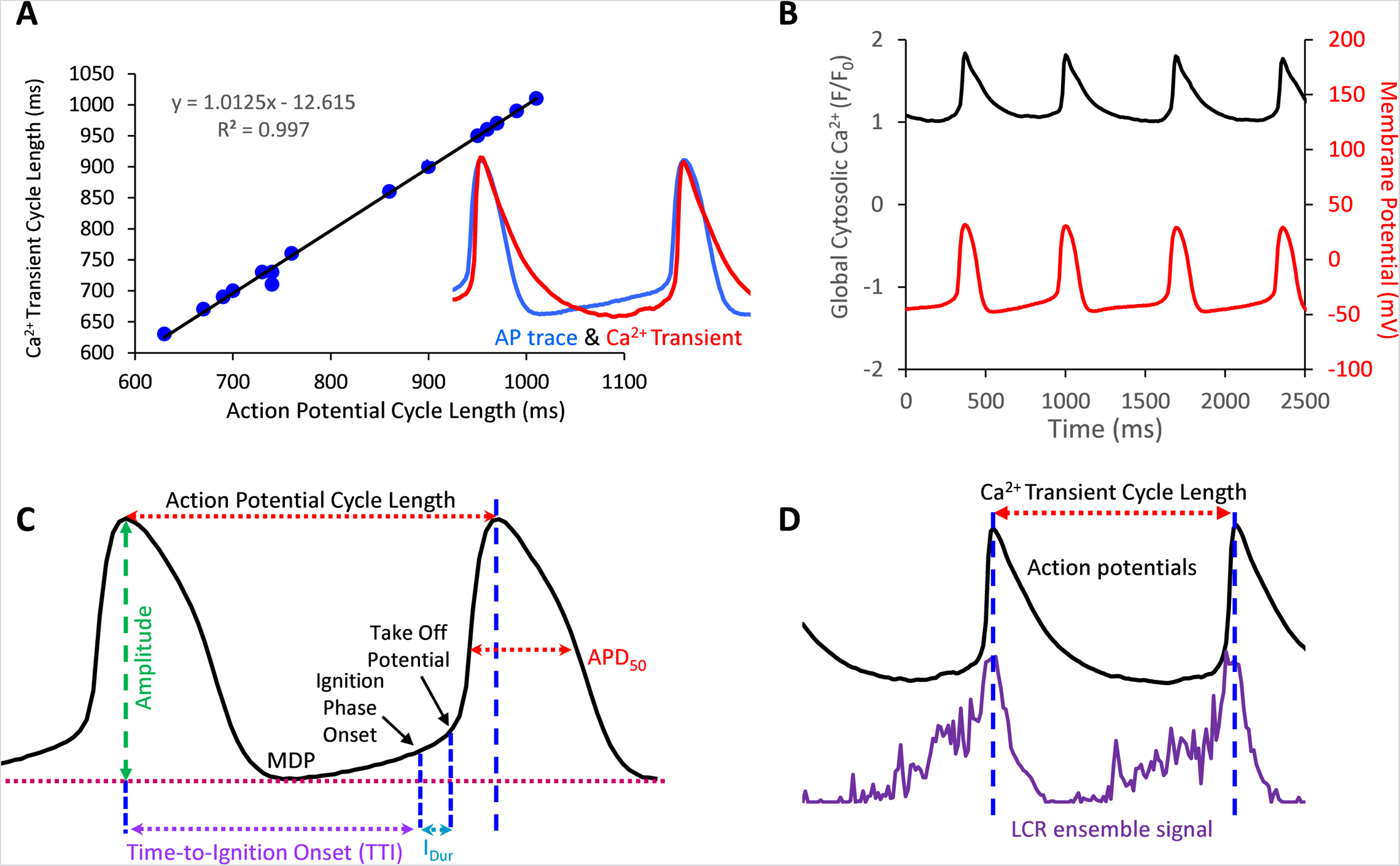
A. Action potentials and AP-Induced Ca^2+^ transients are highly correlated in SANC in which both Ca^2+^ and V_m_ were simultaneously measured (17 cycles measured simultaneously). B, example of a simultaneous V_m_ and Ca^2+^ recording. C and D. Illustrate V_m_ and Ca^2+^ parameters measured in this study.

### Experimental Protocol

In some cells we measured APs only, and in others we measured Ca^2+^ signal only. V_m_ and Ca^2+^ were also recorded simultaneously in a subset of SANC. Both V_m_ and Ca^2+^ recordings were electronically synchronized. The V_m_ measurements and 2D Ca^2+^ signals were obtained before, during, and following washout of 10µM ado (Sigma Aldrich, USA). We measured SANC APCL using two parameters: AP intervals or AP-induced Ca^2+^ transient intervals that were closely correlated (R^2^=0.99, Fig. 1A,B). The rhythmicity of SANC firing was assessed as the coefficient of variation (CV) of AP or Ca^2+^ transient intervals in time series of AP’s or AP-induced Ca^2+^ transient cycles. The mean LCR period was the average of 3-7 LCR periods at baseline and with ado.

### Statistics

Values are expressed as mean ± standard error. Ca^2+^ and electrophysiological measurements in control were compared to those in the presence of ado by one-way ANOVA, paired *t*-test or via Student’s *t*-test, as indicated in the Figure legends. *P* value <0.05 was considered statistically significant.

## Results

### AP Firing Rate and Rhythm

Ado dose-dependently increased AP-induced Ca^2+^ transient cycle length (Fig. 2, Table 1). Ado, at a concentration near its IC_50_ (10µM) (Lou et al., 2014), increased APCL from 492±88 to 687±178 (Table 2). In response to 10µM ado, all rhythmically firing SANC increased APCL, measured via V_m_ or Ca^2+^ signal, and most recovered with washout (Fig. 3). The CV of AP also increased in response to 10 µM ado from 5.1±1.7% to 7.2±2.0% (Table 2). To confirm that the increase in APCL was due to ado effects, time controls of Ca^2+^ and V_m_ measurements were also conducted. There was no significant time effect on either rate of AP firing or AP-induced Ca^2+^ transients’ measurement within thirty minutes (Figure 4).

**Figure 2.**
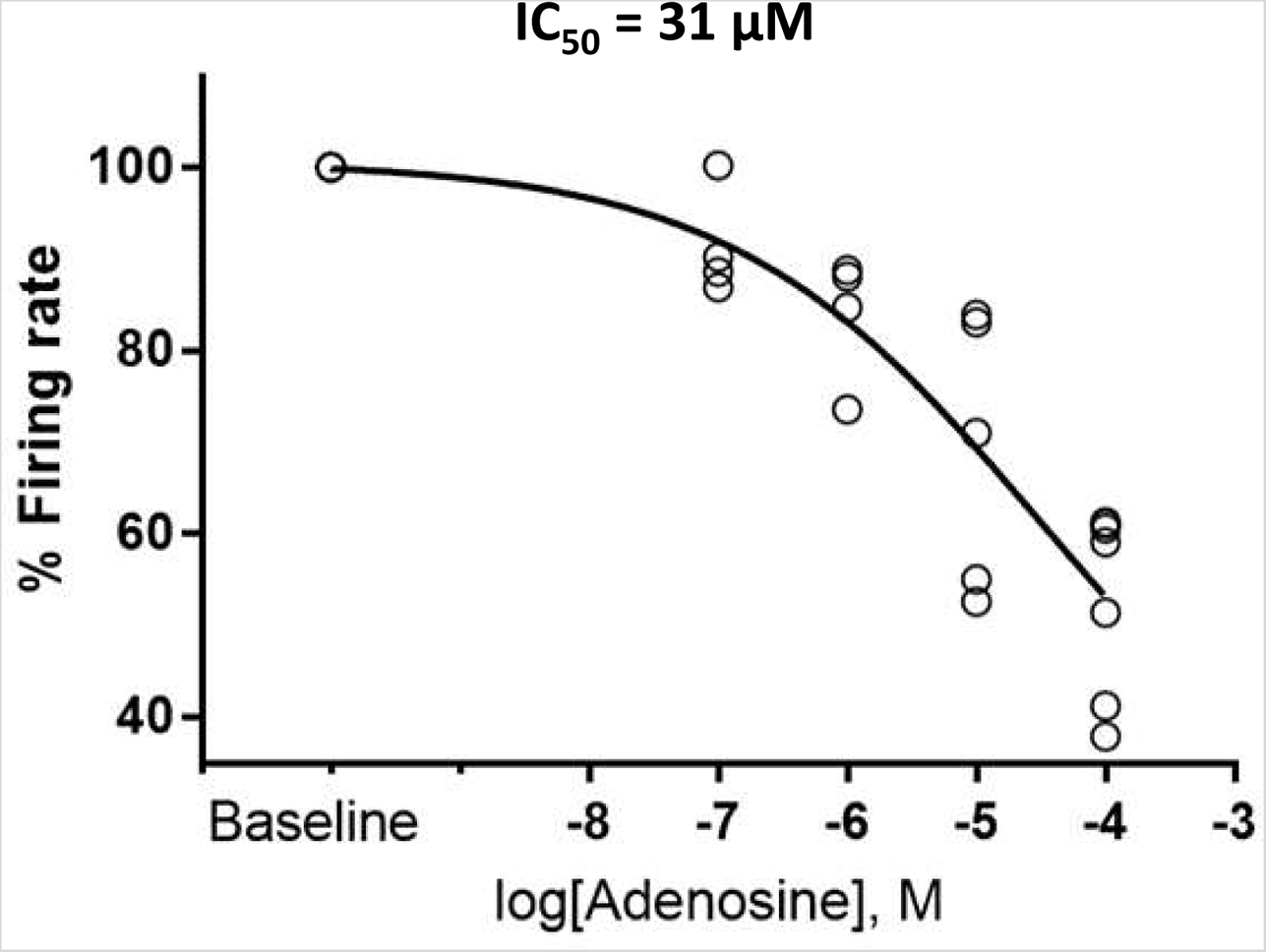
Adenosine slows the firing rate of rhythmically firing SANC in a dose-response manner. The IC_50_ for our dose response curve was 31 µM adenosine. (n=4 for 100nm and 1µM adenosine, n=5 for 10µM adenosine, and n=7 for 100µM adenosine). The data of firing rate response were normalized to be between 0% and 100% of baseline and fitted to Variable slope model using GraphPad program https://www.graphpad.com/guides/prism/latest/curve-fitting/reg_dr_inhibit_normalized_variable.htm. The equation for this fitting function was: Y=100/(1+10^((LogIC50-X)*HillSlope)).

**Figure 3.**
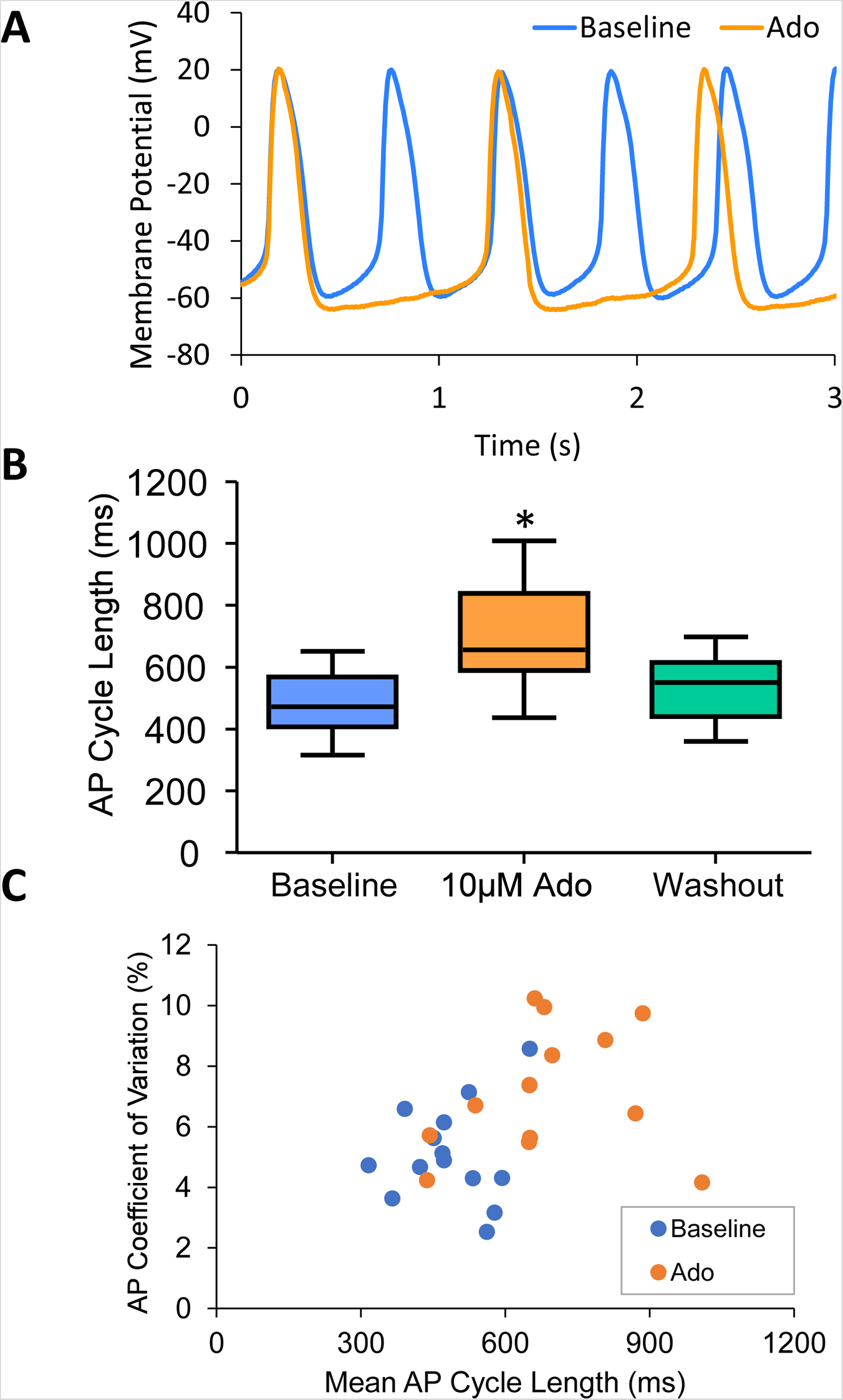
A, an example of a cell that increased APCL from 577ms at baseline 1009ms with adenosine and recovered to 597ms with washout (washout not shown). B, Statistical analysis of adenosine effect: SANC action potential cycle length (APCL) increased with adenosine and recovered with washout in SANC (n=14). C, APCL and coefficient of variation (CV) increased with adenosine. *p<0.05 via one-way repeated measures ANOVA.

**Figure 4.**
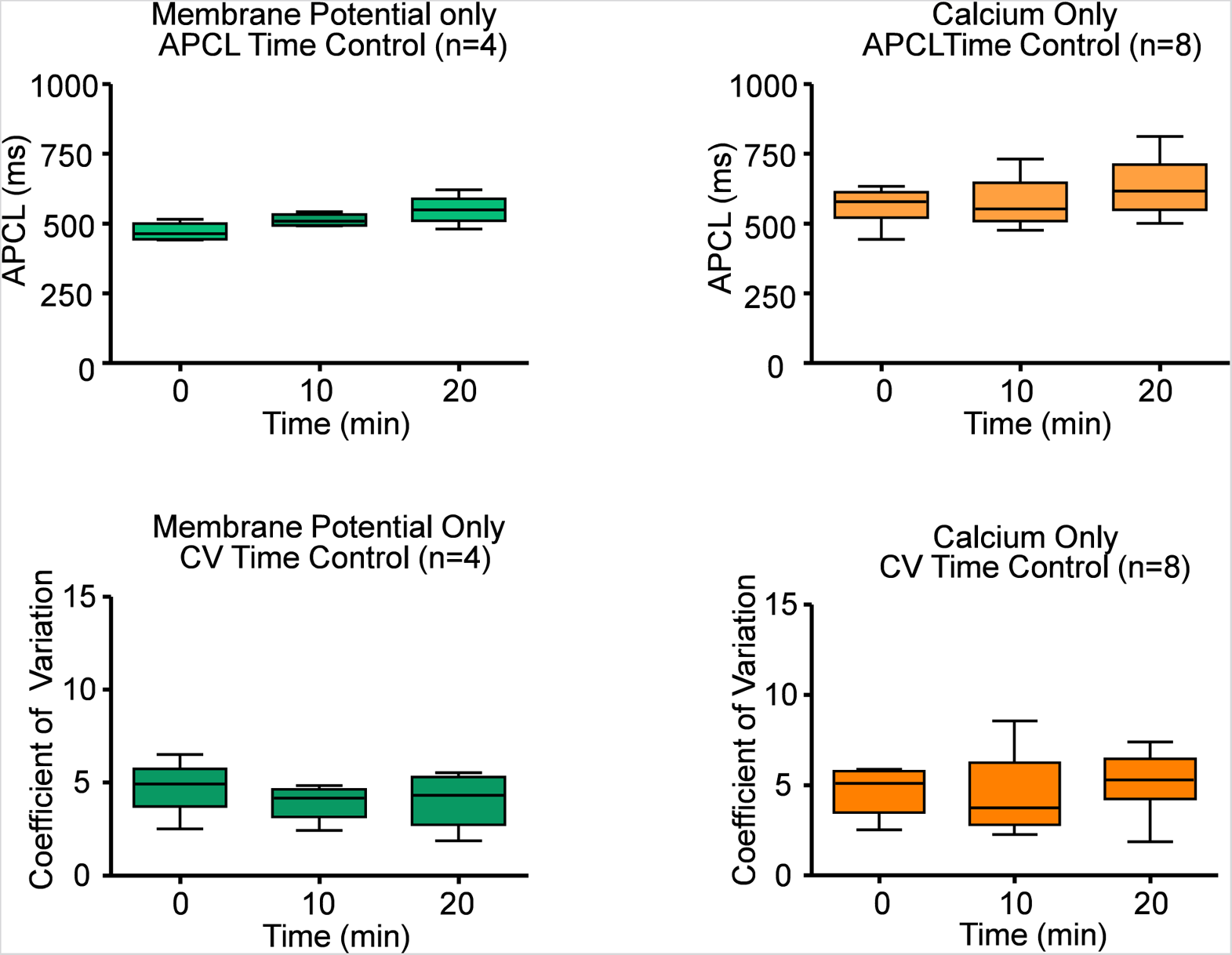
Time controls of Ca^2+^ and electrophysiology measurements show that SANC action potential cycle length (APCL) and coefficient of variation (CV) do not change with time. Repeated measures ANOVAs of Ca^2+^ and V_m_ measurements showed there was no significant time effect on SANC firing rate and CV within twenty minutes. Based on this data, all experiments were performed within twenty minutes.

**Table 1.** Adenosine slows SANC firing rate in a dose-dependent manner. Calcium measurements of rhythmically firing SANC were taken before and after adenosine exposure. To minimize cell phototoxicity, different cells were used for each dose of adenosine. All SANC measured slowed down in response to adenosine. At each concentration of adenosine, the increase in APCL was significant (p<0.05).

| Parameters | Baseline<br>(n=20) | Adenosine<br>100nm<br>(n=4) | Adenosine<br>1μm<br>(n=4) | Adenosine<br>10μm<br>(n=5) | Adenosine<br>100μm<br>(n=7) |
| --- | --- | --- | --- | --- | --- |
| SANC APCL (ms) | 408±70 | 435±43* | 531±65* | 670±277* | 703±222* |
| SANC APCL<br>% Change | 100 | 108±7 | 120±11* | 150±34* | 184±48* |
| Coefficient of<br>Variation (%) | 4.9±2.9 | 7.2±2.8 | 8.9±6.7 | 12.1±12.6 | 10.5±7.9* |

**Table 2.** Membrane potential parameters recorded in this study. Panel A, action potential characteristics of SANC measured before, during, and after adenosine perfusion. The data obtained from 10 single SANCs isolated from five rabbit hearts. *P < 0.05, compared to baseline by two-tailed paired t test. CL; cycle length max-to-max, CV; coefficient of variation, MDP; max diastolic potential, Ampl, amplitude; dVdtmax, maximum upstroke velocity; TTI, time-to-ignition onset; IP, Ignition Potential; lour, Ignition Duration (from IP to TOP); Time to TOP (from MDP to TOP); TOP, take off potential.

| Condition | n | CL (ms) | CV (%) | MDP (mV) | Ampl (mV) | dVdt<sub>max</sub> (V/s) | APD<sub>50</sub> (ms) | Mean DD Slope (mV/s) | TTI (ms) | IP (mV) | I<sub>Dur</sub> (ms) | TOP (mV) |
| --- | --- | --- | --- | --- | --- | --- | --- | --- | --- | --- | --- | --- |
| Baseline | 10 | 492±88 | 5.1±1.7 | -64.1±5.6 | 89.8±9.1 | 3.7±1.1 | 135±28 | 43.0±18 | 478.5±164 | -56.5±8.1 | 24.6±10.5 | -59.0±6.7 |
| Ado | 10 | 687±178* | 7.2±2.0* | -67.7±4.8* | 97.2±9.2* | 3.9±1.4* | 141.6±22.3 | 22.8±9.8* | 773.9±498* | -57.0±7.4 | 28.1±13.6* | -59.4±5.9 |
| Ado,% of baseline | 10 | 143±33* | 142±36* | 105±4.0* | 110±7.8* | 117±14* | 106±15 | 58±25* | 150±38* | 101±3.4 | 112±16 | 101±3.4 |
| Washout | 10 | 554±102 | 5.0±1.2 | -66.1±3.9 | 95.2±9.2* | 3.9±1.2* | 138.6±20.9 | 42.0±29.1 | 518.7±143 | -56.5±8.0 | 26.4±8.7 | -58.5±6.9 |
| Washout % of baseline | 10 | 109±13 | 104±31 | 104±6.9 | 108±9.5* | 115±12* | 107±19 | 100±38 | 111±16 | 100±11 | 109±25 | 101±11 |

### Membrane Potential

Ado prolonged the APCL (Fig. 3A and 3B) confirming prior observations (Belardinelli et al., 1988;Ren et al., 2003). In SANC in which only V_m_ was measured, the ado-induced increase in APCL was washable after 10 minutes (Fig. 3B, Table 2). Ado significantly hyperpolarized maximum diastolic potential (MDP) and significantly increased AP amplitude, maximum upstroke velocity (dVdt_max_), and decreased the mean diastolic depolarization slope (Fig. 3A, Table 2). The significant hyperpolarization of MDP and increases in APCL and dVdt_max_ in response to ado are also consistent with previous studies in isolated SANC (Ren et al., 2003). Concurrently, with the prolongation of mean APCL, APCL variability increased (Fig. 3C, Table 2). Time to Ignition Onset (TTI), defined as the time period from the AP peak to the ignition phase onset (Fig. 1C), shown to predict the APCL (Lyashkov et al., 2018) was also markedly prolonged by ado (Table 1). The ignition process itself was also slowed down evidenced by significantly longer Idur (Table 1).

### Ca^2+^ Transients

Ca^2+^ signals were recorded only from rhythmically firing SANC at baseline (CV<10%). An example of Ca^2+^ signal measurement in a representative cell is shown in Fig. 5A. Ado increased the mean AP-induced Ca^2+^ transient cycle length and its cycle-to-cycle variability (Fig. 5, Table 3) to similar extents as for those measured via perforated patch clamp (Fig. 3, Table 2).

**Figure 5.**
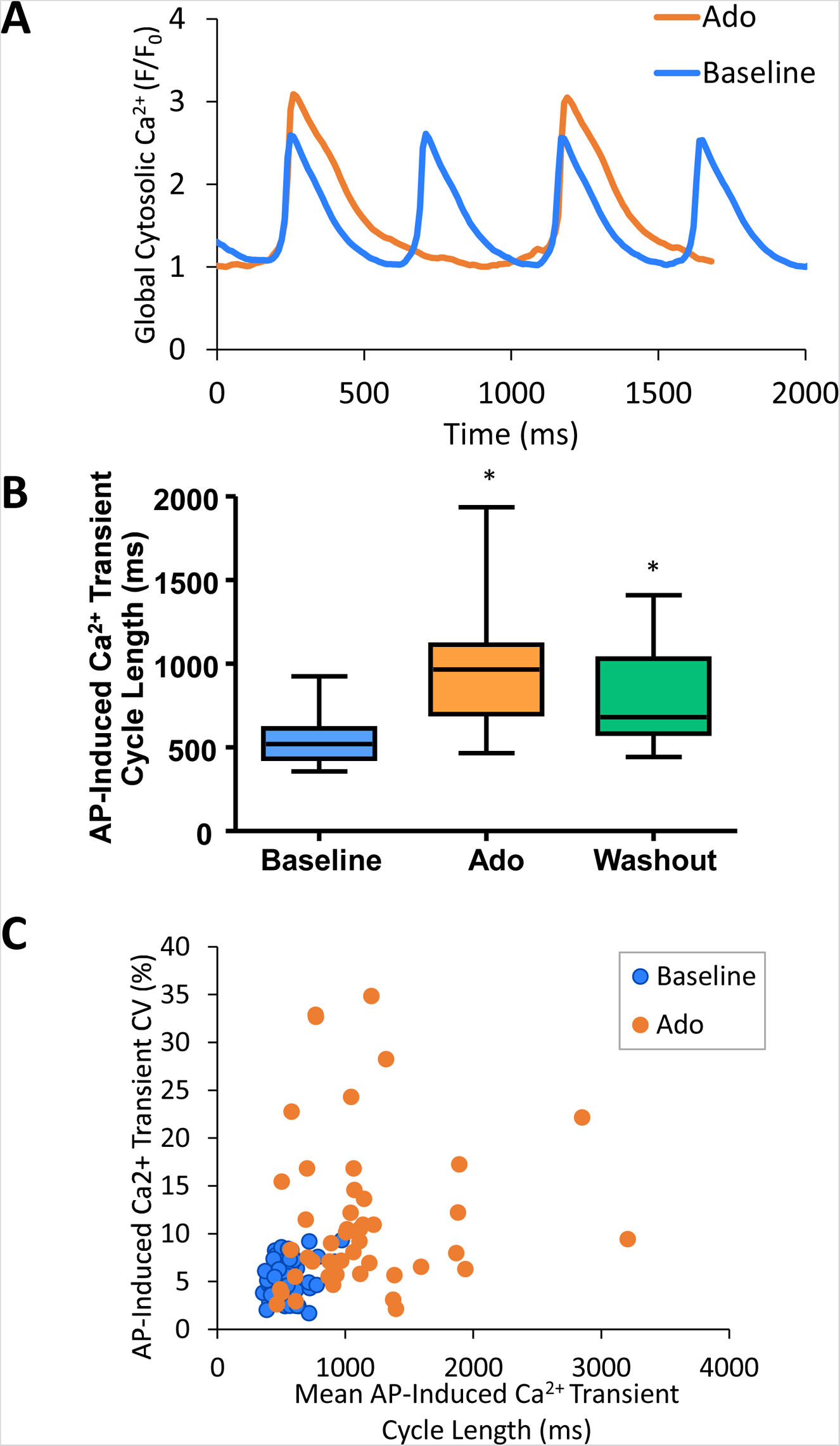
A, an example of a cell that increased APCL from 480ms at baseline 888ms with adenosine and recovered to 648ms with washout (washout not shown). B, SANC AP-induced Ca^2+^ transient cycle length increased with adenosine and recovered partially with washout. (n=46). C, SANC AP-Induced Ca^2+^ transient cycle length and coefficient of variation (CV) increased with adenosine *p<0.05 via one-way repeated measures ANOVA.

**Table 3.** AP-Induced Ca^2+^ transient parameters measured in this study. Panel B, Ca^2+^ measurements in SANC before and during adenosine perfusion (n=40). All cells measured were rhythmically firing at baseline. *P < 0.05, compared to baseline by two-tailed paired t test.

|  | n | CL<br>(ms) | CV (%) | Time to Peak<br>Ca<sup>2+</sup>(ms) | Peak Ca<sup>2+</sup> (F/F<sub>0</sub>) | Time to 90%<br>Decay (ms) |
| --- | --- | --- | --- | --- | --- | --- |
| Baseline | 40 | 540±129 | 5.0±2.2 | 493±115 | 367±252 | 378±114 |
| Adenosine | 40 | 950±331<br>* | 10.6±5.9<br>* | 833±338<br>* | 301±210<br>* | 707±335<br>* |
| Adenosine<br>% of Baseline | 40 | 155±56<br>* | 200±91<br>* | 205±137<br>* | 84±23<br>* | 187±72<br>* |

### Effects of Ado on spontaneous, local diastolic Ca^2+^ releases

The results of statistical analysis of LCR characteristics are presented in Table 4 and illustrated in Fig. 6. Ado did not change the mean number of diastolic LCR events (normalized for longer diastolic times) but reduced the mean LCR size, duration, and ensemble LCR Ca^2+^ signal. When expressed as a percent of control, ado also reduced the LCR size, duration, and ensemble Ca^2+^ signal but also increased the number of LCRs, consistent with the idea that LCRs becomes smaller and less synchronized (i.e. de-synchronized in space and time). With ado, there was an increase in the number and percentage of the smallest LCRs. Correspondingly, there was a decrease in larger LCRs. Ado also prolonged the mean LCR period, consistent with the idea that the period of the coupled-clock system increased.

**Figure 6.**
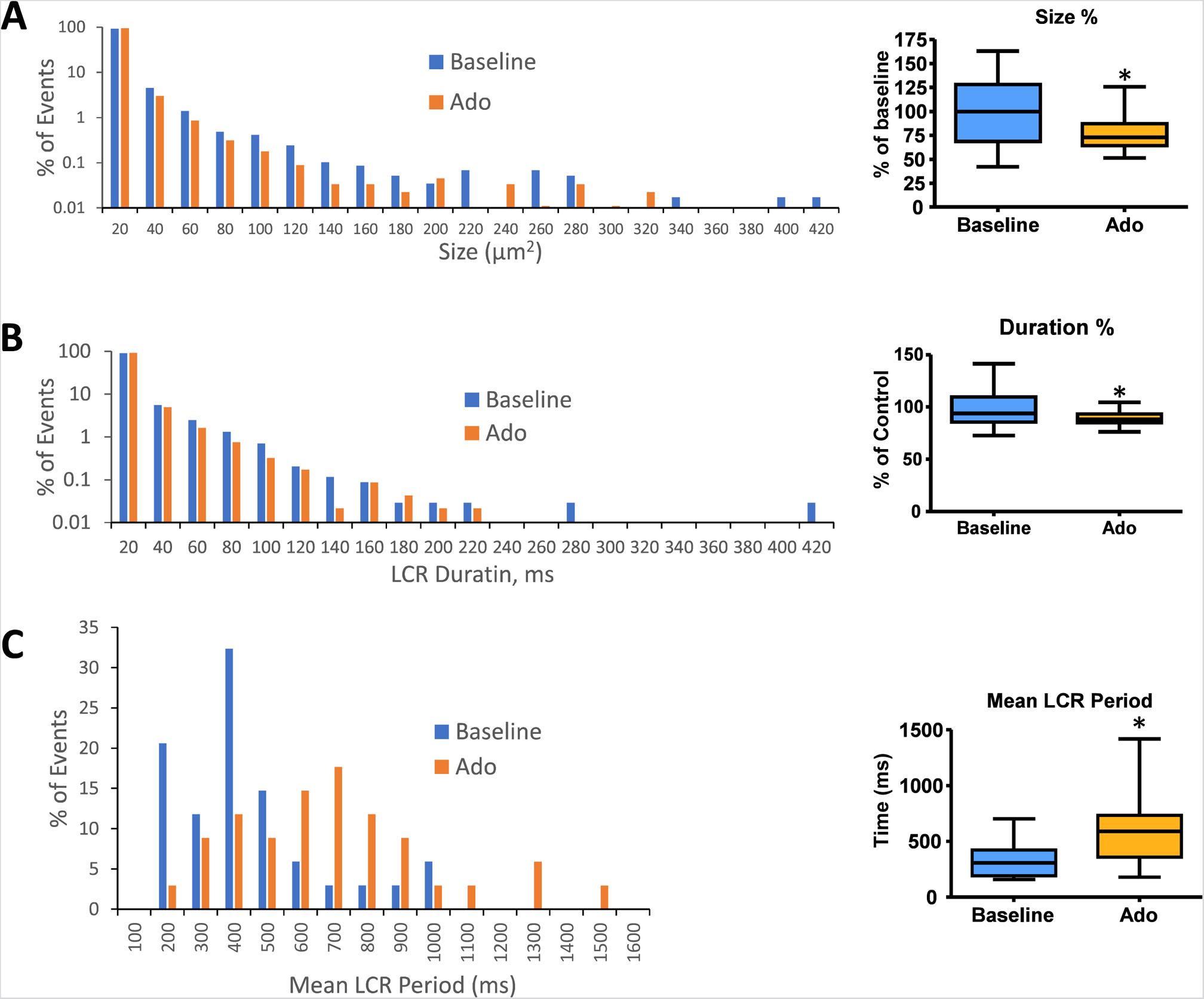
Histograms of LCR size (A) and LCR duration (B), and mean LCR period (C) percentage distributions in all rhythmic SANC before and in the presence of adenosine (n=34). Mean LCR period was the averaged LCR period of 3-7 cycles at baseline and with adenosine for each SANC. Mean LCR period also increased in response to adenosine.

**Table 4.** Local Ca^2+^ release parameters measured in this study. Panel B, Ca^2+^ measurements in SANC before and during adenosine superfusion (n=40). All cells measured were rhythmically firing at baseline. *P < 0.05, compared to baseline by two-tailed paired t test.

| Condition | n | # of LCR Events/<br>Sec | LCR Size<br>($\mu\text{M}^2$) | LCR Duration<br>(ms) | Mean LCR Ensemble<br>($\mu\text{M}^2$) | Mean<br>LCR Period (ms) | LCR Period SD<br>(ms) |
| --- | --- | --- | --- | --- | --- | --- | --- |
| Baseline | 40 | 146±75 | 7.4±2.6 | 16.6±3.4 | 986±437 | 370±179 | 209±107 |
| Ado | 40 | 157±68 | 5.6±2.2* | 14.8±2.6* | 770±358* | 646±320* | 354.8±171* |
| Ado % of<br>Baseline | 40 | 119±42* | 78±19.0* | 89.3±6.5* | 81±22* | 184±83* | 185±88* |

It has been previously shown that a concurrent increase in beat-to-beat variability (CV) accompanied an increase in the mean AP-induced Ca^2+^ transient cycle length and mean LCR period in response to ivabradine, a funny current inhibitor (Yaniv et al., 2014b). The increase in mean AP-induced Ca^2+^ transient cycle length and mean LCR period in response to ado were also accompanied by the increased beat-to-beat variability (CV) of the LCR period and APCL (Fig. 5C, Tables 3 and 4).

### Clock uncoupling in response to Ado

The effect of ado on LCR characteristics resulted in a reduction of the self-organized growth rate of diastolic LCR ensemble Ca^2+^ signal (Fig. 7). At baseline, the majority of SANC showed a strong correlation between LCR period and AP-induced Ca^2+^ transient cycle length, indicating robust clock coupling in most cells (Fig. 7C). In the presence of ado, the mean LCR period and Ca^2+^ transient cycle length increased from baseline but maintained the correlation between LCR period and AP-induced Ca^2+^ transient cycle length (Fig. 7C). For a few cells that deviated from this correlation with ado, as the mean APCL increased in response to ado, a given LCR period was linked to a longer APCL than at baseline (Fig. 7C), reflecting a reduction of the effectiveness of LCR signals to impact the timing of the next AP. This reduced effectiveness manifested as a ‘missed attempt’ at synchronizing between the Ca^2+^ and membrane clock to generate an AP (Fig. 7D). The cells that deviated rightward and upward the most from the correlation between LCR period and AP-induced Ca^2+^ transient cycle length with ado experienced the most uncoupling.

**Figure 7.**
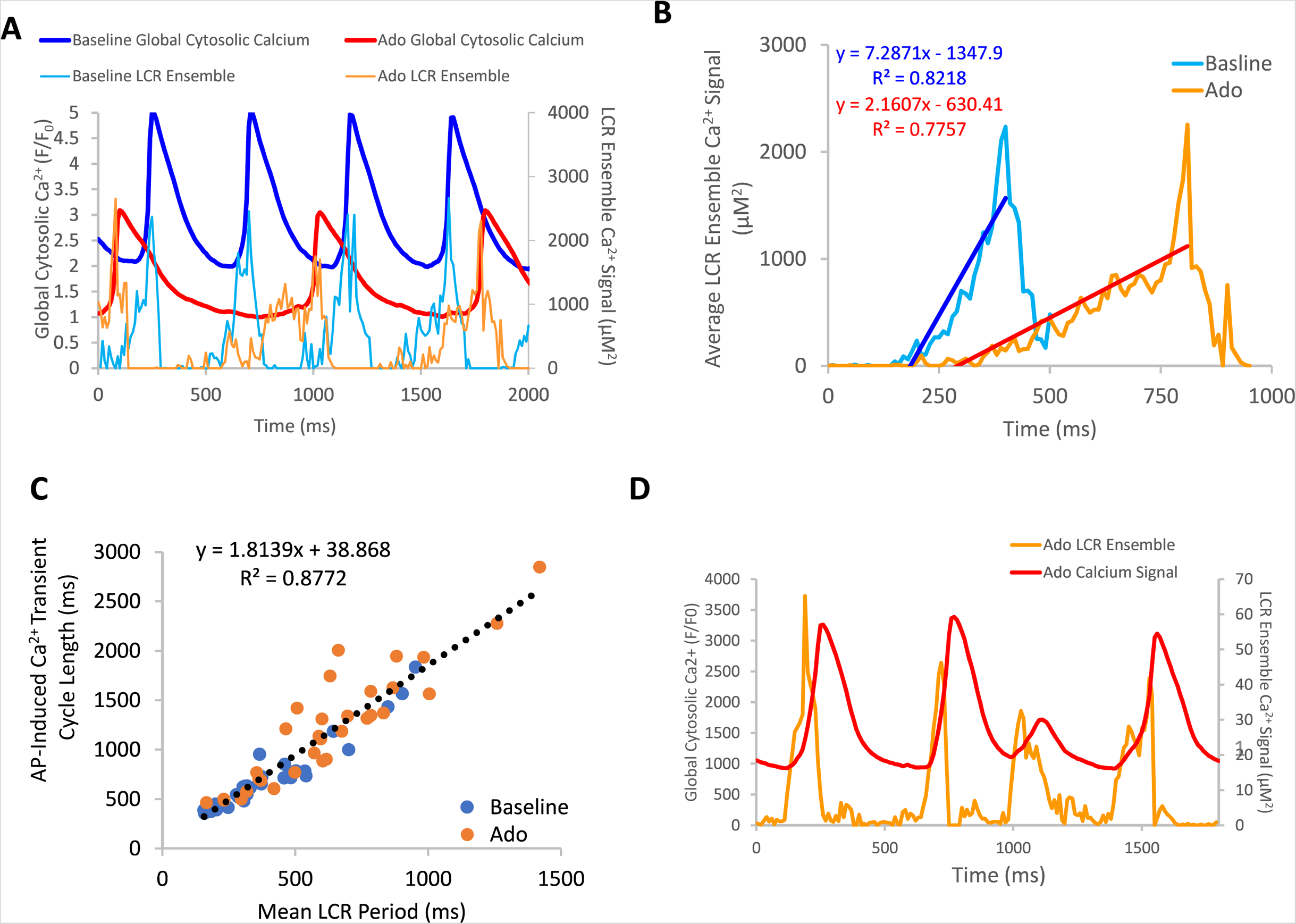
A, an example of a rhythmically firing SANC that decreases in rate of LCR ensemble growth in response to adenosine. There is an overall decrease in global cytosolic Ca^2+^ with adenosine compared to baseline. Panel B depicts the average LCR ensemble growth rate of 5 cycles at baseline and with ado. The average LCR ensemble growth rate decreased to 42% of baseline with adenosine. This decrease reflects the changes in intracellular Ca^2+^ availability, manifesting in changes to LCR parameters. This results in an extended time period of Ca^2+^ cycling between AP-induced Ca^2+^ transients. Panel C, the correlation between mean LCR Period (3-7 beats) and AP-induced Ca^2+^ transient cycle length is reduced at longer cycle lengths. At baseline, there is a strong correlation between APCL and LCR period, indicating robust clock coupling (n=35). Adenosine, in the same population of cells, the mean APCL and LCR period increased for many cells. For a subset of cells, mean APCL increased more than mean LCR period, indicating reduced fidelity of clock coupling indicated by the orange circles that digress up and leftward from the linear trendline (dashed line). The linear trendline includes baseline and ado values. Panel D, an example of a SANC with ado where LCR ensemble growth propagation was insufficient and failed to generate an AP, resulting in clock uncoupling and a ‘failed AP attempt’.

### Simultaneous measurements of membrane potential and Ca^2+^

The aforementioned measurements of Ca^2+^ and V_m_ were made in different cells. To directly assess the effect of ado on clock coupling, simultaneous membrane and Ca^2+^ measurements were performed within the same cell prior to and following ado superfusion (Movie S1, Fig. 8). The prolongation and increased variability of APCL in the presence of ado occurred concurrently with a reduced growth rate of the LCR ensemble Ca^2+^ signal (Fig. 9, Panel B). The AP ignition times, LCR periods, and APCLs of five cells in which V_m_ and Ca^2+^ were measured simultaneously are illustrated in Fig. 9. The relationship between TTI and APCL in these five cells is illustrated in Fig. 9A. Note how TTI and APCL increase in the presence of ado and maintain their relationship. TTI informs on the LCR period, even for extremely uncoupled cells (r^2^=0.9) (Fig. 9C) because the growth of the ensemble LCR Ca^2+^ signal, via its effect to increase more inward I_NCX_, initiates the ignition phase recorded as TTI (Lyashkov et al., 2018). Note a concurrent shift in TTI and LCR period at baseline and in the presence of ado (Fig. 9C). Therefore, the APCL depends on the LCR period (Fig. 9B) and that relationship informs on the fidelity of clock coupling: with ado, the clock coupling, reflected in the LCR period, was reduced.

**Figure 8.**
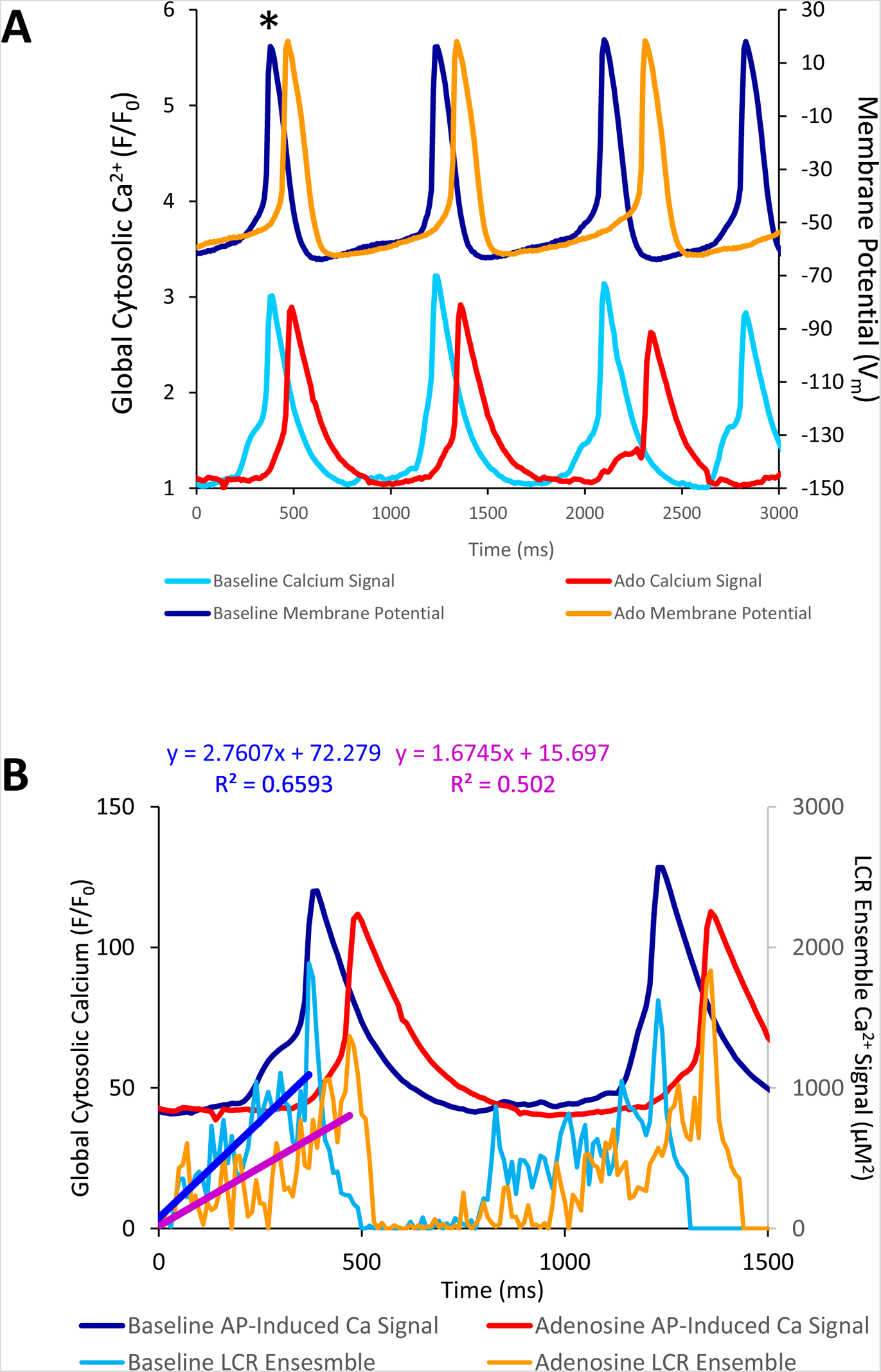
A, simultaneous V_m_ and Ca^2+^ recordings of a rhythmically firing SANC at baseline that fires with decreased rhythmicity in response to 10 µM Ado. B, the rate of LCR ensemble growth is steeper at baseline (blue line) and decreases in response to adenosine (magenta line). The asterisk indicates the baseline and ado cycle lengths measured in Figure 8A.

**Figure 9.**
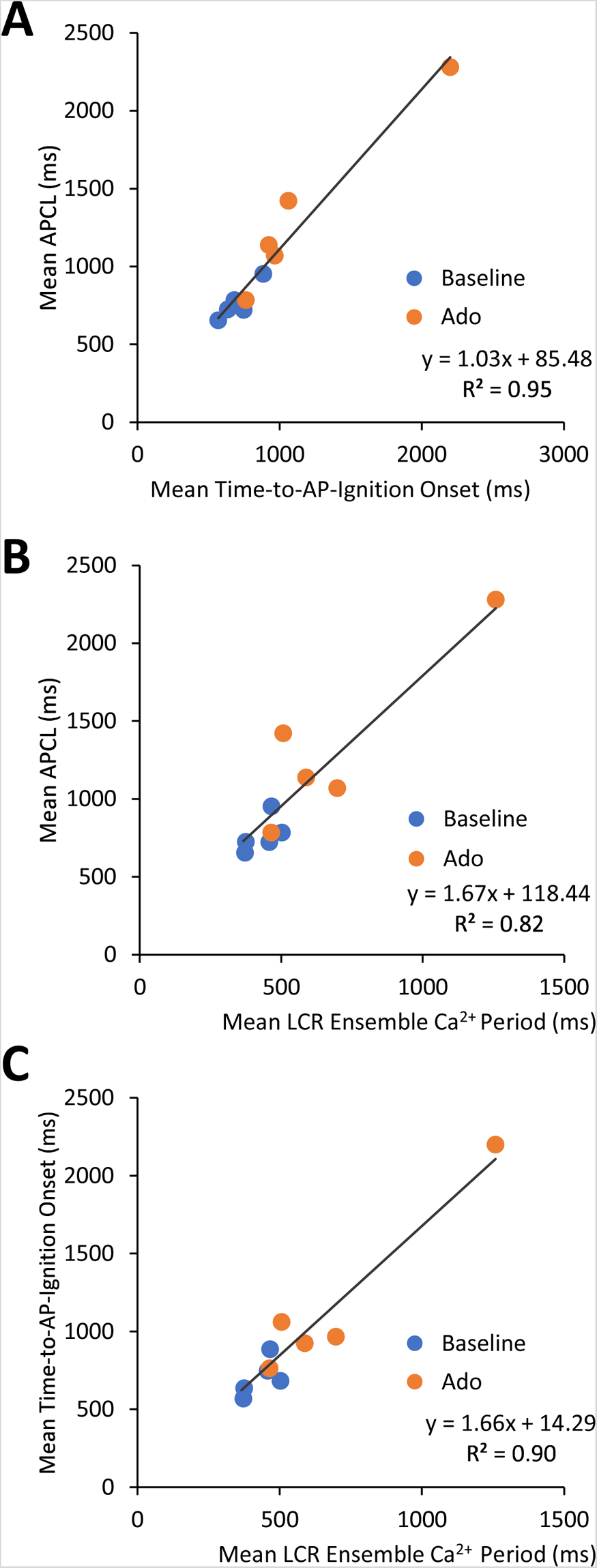
Relationships between key parameters of the coupled-clock system in control (baseline) and in the presence of adenosine in SANC in which Ca^2+^ and V_m_ were measured simultaneously. The LCR period informs on the APCL because the LCR period informs on the time-to-AP-ignition onset. Both LCR period and Time-to-AP-ignition onset increase with adenosine.

### V_m_-Ca^2+^ Phase-Plane Diagrams

Phase-plane diagrams of V_m_ versus Ca^2+^ permit closer inspection of simultaneous time-dependent changes occurred in V_m_ and Ca^2+^. V_m_ and Ca^2+^ throughout the AP cycle inform on the electrochemical gradient oscillation that underlies each AP cycle (Lakatta et al., 2003). The phase-plane diagrams of the AP and Ca^2+^ recordings prior to and during ado superfusion are illustrated in Fig. 10. Effects of ado on both clock parameters (Figs 3 and 5, Tables 2, 3, and 4) would be expected to alter the electrochemical gradient oscillation characteristics exemplified by the V_m_-Ca^2+^ diagram.

**Figure 10.**
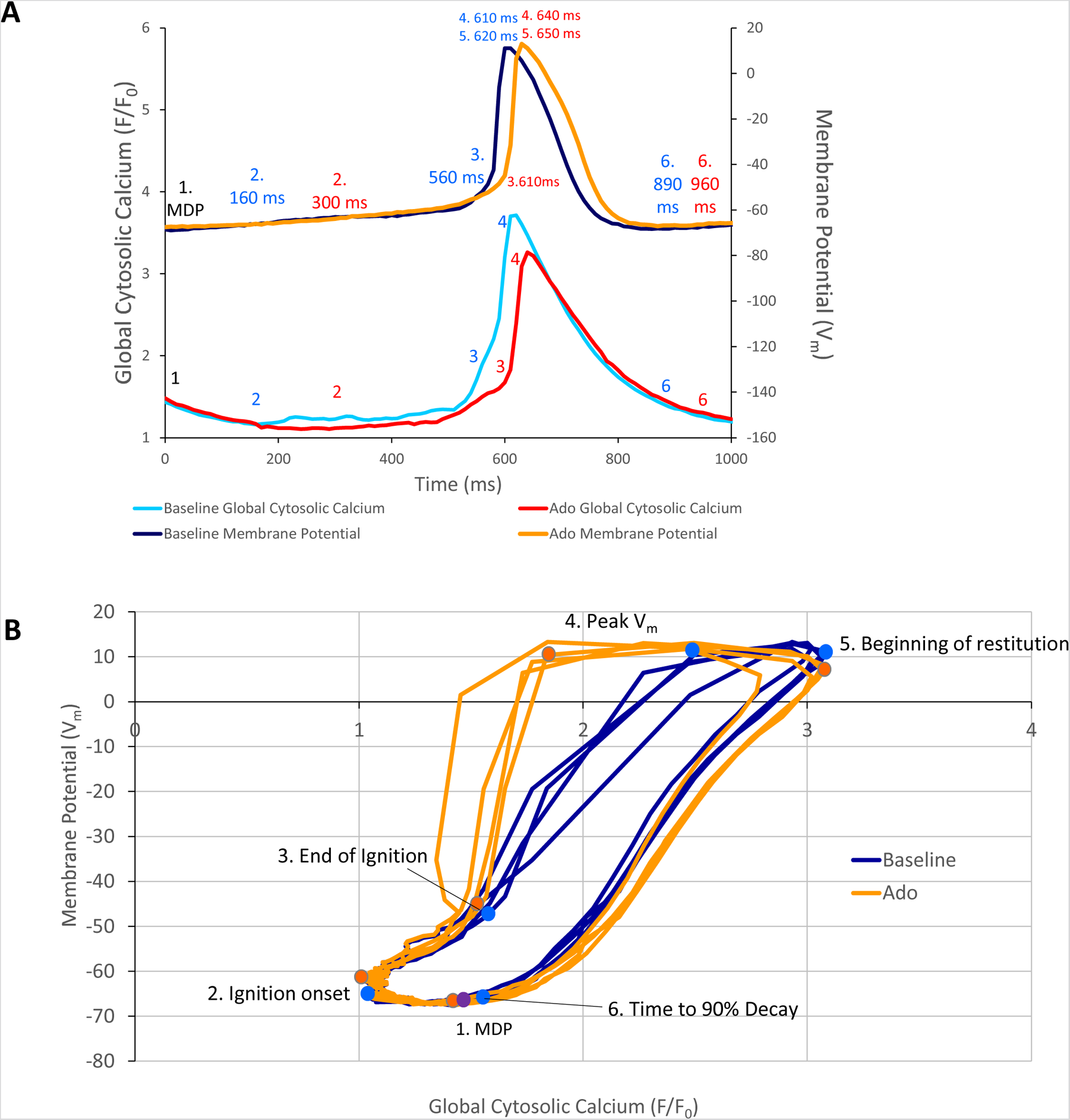
A, Phase V_m_-Ca^2+^ relationship. A, one AP cycle (* in figure 5) prior to (baseline) and during adenosine (Ado) superfusion in which V_m_ and Ca^2+^ were measured simultaneously. B, phase V_m_-Ca^2+^ diagram depicting the relationship between V_m_ and global cytosolic Ca^2+^ during several AP cycles (electrochemical gradient oscillations) at baseline (blue) and with adenosine (orange); 4 consecutive cycles from Figure 5 are shown. Numbers 1-5 indicate the times during the AP cycles in Panel A.

Point 1 in Fig. 10 Panels A and B marks the MDP in control and ado. The time from the MDP to ignition onset (labeled 2 in Panels A and B) occurred at a similar V_m_ and lower global Ca^2+^ in the presence of ado than at baseline. The AP ignition phase onset occurred earlier at baseline (160 msec) than in the same cells in the presence of ado (300 msec) (Fig. 10, Panel A). Progressive Ca^2+^ ensemble self-organization and its effect on V_m_ and on the global Ca^2+^ signal caused the ignition process to proceed from 2 to 3, the take off potential of the AP, in Panels A and B. The take off potential marks the end of the ignition phase (labeled 3 in Panels A and B). The time from ignition onset to TOP (take off potential) is the duration of the ignition phase. Ado prolonged the time-to-ignition (MDP to ignition phase onset) from 160msec at baseline to 300msec with ado. The ignition phase duration decreased with ado from 400 msec at baseline to 310 msec with ado Pane A). Peak V_m_ and Ca^2+^ transient amplitudes (labeled 4 in Panels A and B) occur at later times with ado (640 msec) than baseline (610msec) (Panel A). AP repolarization and Ca^2+^ transient decay initiates sooner at baseline than during ado (Panel A). Peak V_m_ amplitude, labeled 4 in Panels A and B, occurred at a lower Ca^2+^ level but later in time in ado (640msec) than baseline (610msec) (Fig. 10A and B). The time to 90% restitution were 890msec at baseline and 960msec with ado at Point 6 (Panel A and B).

Note that the degree of hysteresis indicates the overall uncoupling of Ca^2+^ and V_m_ signals during an AP cycle. This hysteresis is greater with ado than at baseline. Whereas the difference of the duration of the AP ignition period is 560 to 610 msec (1.1 times greater) and time to 90% decay is 890msec and 960msec (1.08 times greater), the main difference to the time domain and percent differences between ado and baseline is the time-to-ignition onset. The major factor increasing hysteresis between ado and control in the phase-plane diagram is due to a 2-fold increase in the time-to-ignition onset in ado (300 msec) and control (160 msec). The time-to-ignition onset is regulated by the pumping of Ca^2+^ into the SR. This rate of Ca^2+^ cycling into the SR to achieve the threshold required for spontaneous LCRs to occur and self-organize, is the degree to which it influences V_m_ time-to-ignition onset.

## Discussion

The present study builds upon the known effects of ado to slow SANC AP firing and change V_m_ characteristics (Belardinelli et al., 1988). However, more recent studies have demonstrated that it is a coupled membrane and Ca^2+^ clock system that regulates SANC AP firing rate and rhythm. Ca^2+^ is an important oscillatory substrate involved in crosstalk between surface membrane and Ca^2+^ clocks (Fig. 11). The availability of intracellular Ca^2+^ is regulated by the balance of Ca^2+^ influx and efflux from the cell, which decrease in response to decreased clock protein phosphorylation (Fig. 12). The present study is the first to measure and demonstrate that ado reduces and slows down intracellular Ca^2+^ cycling by reducing mean LCR size and duration and increased mean LCR period. The fidelity of clock coupling stems from the tight relationship between Ca^2+^ cycling and V_m_. Thus, ado modulates Ca^2+^ and membrane clocks, resulting in ado-induced clock uncoupling.

**Figure 11.**
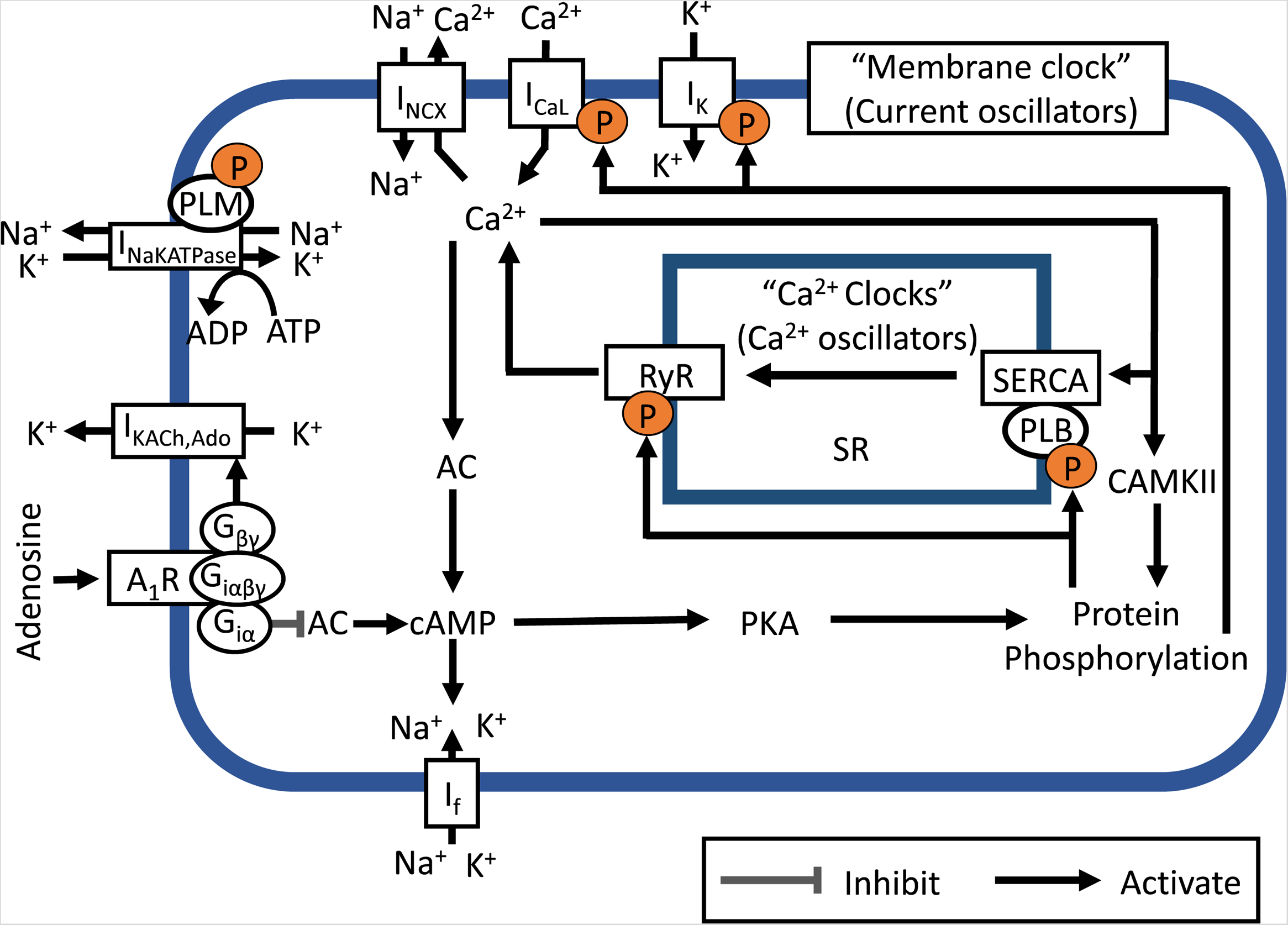
Simplified schematic of the coupled system of membrane ion current oscillators and Ca^2+^ oscillators (Coupled-clock system) operative within sinoatrial nodal cells (SANC). The system provides robust and flexible AP firing rates. Constitutively active Ca^2+^-AC-cAMP-PKA signaling intrinsic to SANC, that is activated by Ca^2+^ oscillations, couples to an ensemble of electrogenic surface membrane molecules (current oscillators). AC, adenylyl cyclase; cAMP, cyclic AMP; PKA, protein kinase A; A_1_R, adenosine A_1_ receptor; GI_α,_ GI protein alpha subunit; G_βγ,_ GI protein beta gamma subunit; RyR, ryanodine receptor; SERCA, sarco-endoplasmic reticulum Ca^2+^-ATPase; PLB, phospholamban; SR, Sarcoplasmic reticulum; CaMKII, Ca^2+^/calmodulin-dependent protein kinase II; I_K_, delayed rectifier K^+^ current; I_KACh,Ado_, Acetylcholine/Adenosine-activated K^+^ current; I_f_, HCN currents; I_CaL_, L-type Ca^2+^ current; I_NCX_, Na^+^/Ca^2+^ currents; and I_NaKATPase_, Na^+^/K^+^ pump.

**Figure 12.**
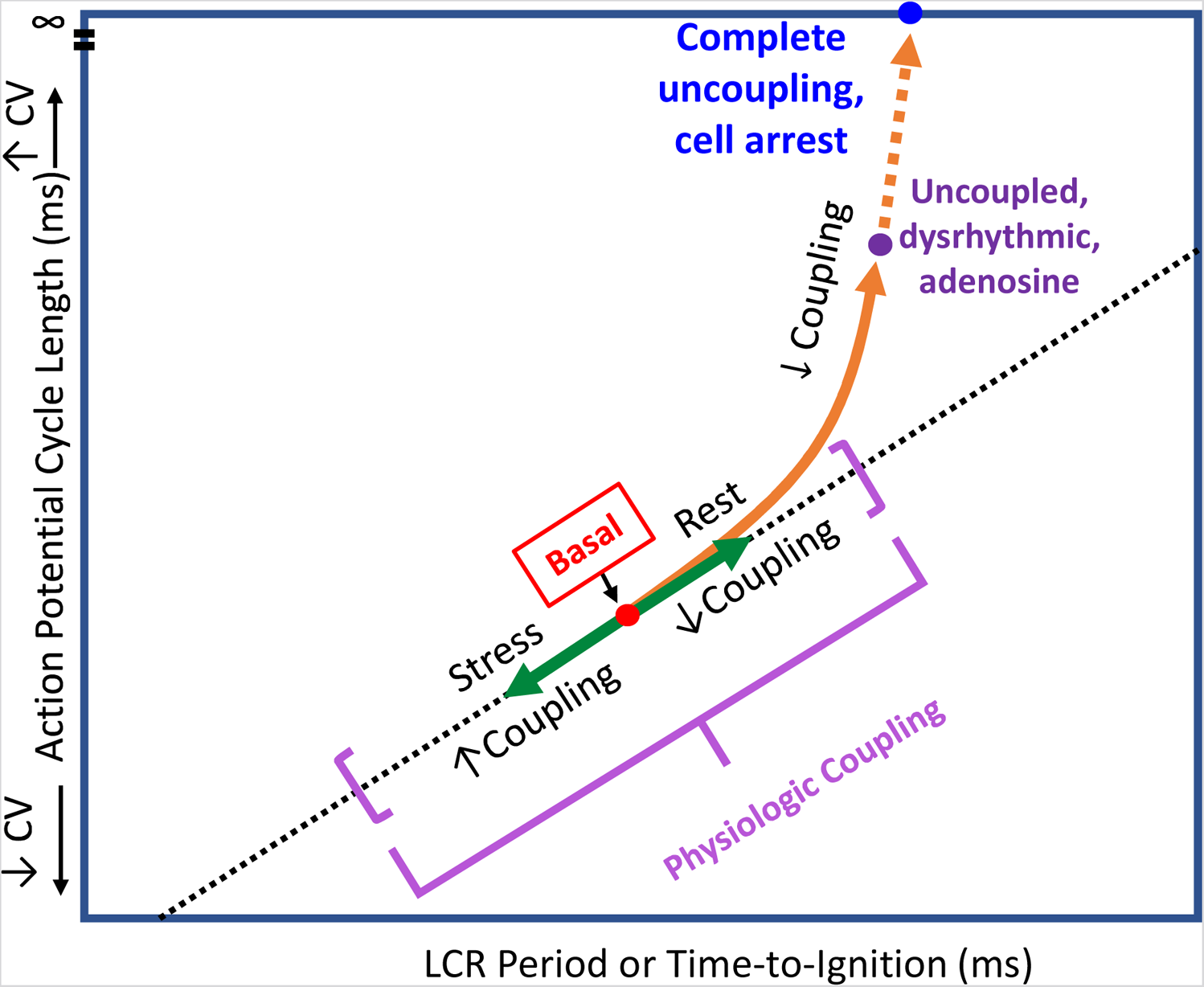
A schematic for SANC clock coupling/uncoupling. Clocks are coupled in the awake basal state, in response to physiologic stress (beta-adrenergic stimulation) coupling increases and beat-to-beat variability is reduced. In physiologic responses to vagal stimulation or adenosine, the clocks become partially uncoupled and the APCL becomes prolonged. These effects become more exaggerated as Gi coupled stimulation increases at higher drug concentrations.

The present study demonstrates that ado increases APCL and APCL variability of SANC by directly and indirectly effecting both membrane and Ca^2+^ clocks. Ado directly activates I_KACh, Ado_ to hyperpolarize SANC and slow firing rate, thus directly effecting the membrane clock. Ado also has indirect effects on the membrane clock via G_iα_ (Fig. 11) to reduce intracellular protein phosphorylation and SR Ca^2+^ cycling to further slow SANC firing rate. Ado exerts direct effects on the Ca^2+^ clock by decreasing intracellular phosphorylation of Ca^2+^signaling proteins (like Ach). Ado indirectly effects the Ca^2+^ clock via its effect on the membrane clock to slow SANC firing rate, commensurate a decrease in net SR Ca^2+^ influx and intracellular Ca^2+^, i.e. oscillatory substrate for Ca^2+^clock. Thus, changes in SANC firing rate were due to direct effects on both clocks and indirect effects of both clocks on each other. Numerous feedbacks and feedforwards of this process occurred until a new equilibrium was reached.

Mechanisms by which ado slows AP firing and increases beat-to-beat variability are similar to cholinergic signaling via Ach. Ach and ado have different membrane receptors, but likely target the same I_KAch_ channels via G_βγ,_ hyperpolarizing the cell membrane and extending the time of diastolic depolarization (Kurachi et al., 1986). Ach, like ado, activates G_iα_ to inhibit AC activity and reduce cAMP-mediated, PKA dependent phosphorylation of downstream Ca^2+^ cycling proteins targets (DiFrancesco and Tromba, 1988;Dessauer et al., 1996;Lyashkov et al., 2009). While the present study did not directly measure ado’s effects on phosphorylation of clock proteins, it has been previously demonstrated that cholinergic receptor stimulation, like ado, decreases intracellular cAMP and phosphorylation via inhibition I_KACh, Ado_ (Lyashkov et al., 2009). Since ado acts via the same signaling pathway, its effects on intracellular Ca^2+^ cycling observed in this study likely result from the same changes to intracellular phosphorylation.

Other studies have demonstrated that because the membrane and Ca^2+^ clocks are tightly coupled, specific inhibition of a specific molecule in one clock that reduces SANC AP firing rate indirectly affects the function of the other clock and affects clock coupling fidelity. For example, specific I_f_ inhibition by ivabradine not only reduces SANC firing rate and increases AP cycle variability, but also indirectly reduces intracellular Ca^2+^ (Yaniv et al., 2014a). Thus, the overall Ca^2+^ effect of ivabradine, and likely ado, to reduce AP firing rate involves effects on both clocks as well as clock coupling.

Cyclopiazonic acid (CPA) is a specific Ca^2+^ clock inhibitor that selectively and reversibly inhibits SERCA Ca^2+^-ATPase (Goeger et al., 1988;Nelson et al., 1994). CPA been shown to dose-dependently decrease SANC firing rate by suppressing SERCA-mediated Ca^2+^ pumping (Vinogradova et al., 2010). The slowing of SANC firing with CPA was reflected by decreased SANC LCR size and number and increased LCR period (Vinogradova et al., 2010). The CPA-induced changes in LCR characteristics delayed the occurrence of LCR activated Na^+^-Ca^2+^ current and reduced its amplitude, contributing to a decreased mean diastolic depolarization slope (Vinogradova et al., 2010). Thus, while CPA directly modified only Ca^2+^ clock function via the suppression of SERCA function, the collective effect on M and Ca^2+^ clock characteristics resulted in the APCL prolongation of the SANC firing. These results can be interpreted to indicate that any disturbances of the Ca^2+^ clock by ado in our experiments would also influence both clocks and their coupling, ultimately reducing SANC firing rate.

Adenosine is known to be released in response to metabolic stress such as hypoxia and inflammation (Grenz et al., 2011;Idzko et al., 2014). Its effect on clock coupling in vitro may explain its function in vivo. Upregulation of A_1_R protein expression and increased plasma levels of ado are also found to be associated with heart failure and ischemia (Newman et al., 1984;Funaya et al., 1997;Lou et al., 2014). Evidence is accumulating that adenosine contributes to SAN dysfunction in heart failure (Lou et al., 2014). New insights, through 2D optical mapping of the intact SAN, have shown that failing SAN is increasingly sensitive to ado (Lou et al., 2014;Li et al., 2017). This increased influence of ado has been found to contribute to SAN dysfunction by amplifying intrinsic conduction abnormalities such as atrial fibrillation and sinus exit block (Li et al., 2017). Given the significant contribution of adenosine to SAN dysfunction, its effect on isolated SANC in vitro shown here may have some baring on arrhythmias in the context of heart failure and ischemia.

## STUDY LIMITATIONS

The AP-induced Ca^2+^ transient cycle length was, on average, 13% higher in cells loaded with Ca^2+^ indicator Fluo-4 than cells not loaded with Fluo-4. This may reflect Ca^2+^ indicator buffering of SANC measured via Ca^2+^ transients. Nevertheless, ado effects on mean AP-induced Ca^2+^ transient and APCL variability were similar to the effects of ado on APCL and APCL variability recorded via perforated patch clamp in the absence of Ca^2+^ indicator.

## DATA AVAILABILITY STATEMENT

The raw data supporting the conclusions of this article will be made available by the authors, without undue reservation.

## ETHICS STATEMENT

The animal study was reviewed and approved by the Animal Care and Use Committee of the National Institutes of Health.

## AUTHOR CONTRIBUTIONS

ANW and KT: performed the experiments and analyzed the data. ANW and EGL: drafted the manuscript. ANW, VAM, EGL: critically revised the manuscript for important intellectual content, analysis, and interpretation of the data. VM and EGL: project planning, analysis and interpretation of the data, conception and design of the experiments. All authors approved the submitted version.

## SUPPLEMENTARY MATERIAL

The Supplementary Material for this article can be found online at https://www.frontiersin.org

It includes supplementary Movie S1. The movie includes two videos of Ca^2+^ signals recorded at baseline and in the presence of ado and respective plots of time series of whole-cell cytosolic Ca^2+^, Membrane potential, and LCR ensemble signal.

